# Hippocampal theta coordinates memory processing during visual exploration

**DOI:** 10.1101/629451

**Authors:** James E. Kragel, Stephen VanHaerents, Jessica W. Templer, Stephan Schuele, Joshua M. Rosenow, Aneesha S. Nilakantan, Donna J. Bridge

## Abstract

The hippocampus supports memory encoding and retrieval, with distinct phases of theta oscillations modulating the amplitude of gamma-band activity during each process. Encoding and retrieval operations dynamically interact over rapid timescales, especially when sensory information conflicts with memory. The ability to link hippocampal dynamics to specific memory-guided behaviors has been limited by experiments that lack the temporal resolution to segregate when encoding and retrieval occur. To resolve this issue, we simultaneously tracked eye movements and hippocampal field potentials while neurosurgical patients performed a spatial memory task. Novelty-driven fixations increased phase-locking to the theta rhythm, which predicted successful memory performance. Theta to gamma phase amplitude coupling increased during these viewing behaviors and predicted forgetting of conflicting memories. In contrast, theta phase-locking preceded fixations initiated by memory retrieval, indicating that the hippocampus coordinates memory-guided eye movements. These findings suggest that theta oscillations in the hippocampus support learning through two interleaved processes: strengthening the encoding of novel information and guiding exploration based on prior experience.

## Introduction

Hippocampal theta rhythms are prominent during active exploration of novel environments, perhaps due to encoding and retrieval processes necessary to guide ongoing behavior. Interactions between encoding and retrieval support many important functions, including memory updating, which requires comparing novel sensory inputs to prior memories and integrating the new content into the memory representation (Bridge and Paller, 2012; Bridge and Voss, 2014b). Retrieval-mediated reconsolidation requires the presence of novel information during reactivation, suggesting that this hippocampal-dependent learning process is sensitive to mismatch (associative novelty) between the retrieved content and sensory input (Morris et al., 2006; Winters et al., 2011). Some studies have even demonstrated hippocampal involvement in associative novelty (Bridge and Voss, 2014a; Chen et al., 2013; Duncan et al., 2009; Duncan et al., 2012; Honey et al., 1998; Howard et al., 2011; Kumaran and Maguire, 2007a, 2009; Long et al., 2016; Thakral et al., 2015). However, the underlying novelty and retrieval processes have not been segregated as they unfold, in part because it is difficult to segregate these mechanisms in real time, as they interact continuously during learning. Many experimental designs capitalize on an artificial separation between encoding and retrieval phases, but these designs do not capture the natural interplay between these states that guides exploratory behavior and informs decision-making. Here, we assayed the engagement of encoding and retrieval processing in real time by designing a task to link memory-guided eye movements to intracranial recordings of hippocampal activity, and aimed to identify how theta oscillations are distinctly involved in encoding and retrieval processes in real time.

Eye movements provide rich temporal information regarding the focus of attention and the specific cognitive processes engaged at any given moment (Bridge et al., 2017; Bridge and Voss, 2014b, 2015; Voss et al., 2011). In human and non-human primates, learning through exploration heavily depends on the visual system (Meister and Buffalo, 2016), with eye movements resetting the phase of theta during learning of novel visual information (Hoffman et al., 2013; Jutras et al., 2013). Eye movements are also deployed rapidly, with median saccade rates of 3-5 Hz during visual exploration tasks (Wilming et al., 2017). Thus, eye movements provide an ideal behavioral measure to dissect learning processes on a moment-to-moment basis.

The interaction of fast gamma-band and theta oscillations in the hippocampus could play a key role in coordinating interactions between encoding and retrieval during exploration. The amplitude of gamma increases at specific phases of the theta cycle to support memory processing (Axmacher et al., 2010; Lega et al., 2016). Computational models of hippocampal function suggest that gamma activity associated with encoding and retrieval preferentially occurs at the trough and peak of the theta rhythm, respectively (Hasselmo et al., 2002). Task-based observations of hippocampal firing support this idea, showing that novel stimulus encoding and memory retrieval are enhanced at distinct phases of the theta cycle (Douchamps et al., 2013; Lever et al., 2010; Manns et al., 2007; Newman et al., 2013). In addition, closed-loop optogenetic stimulation of inhibitory neurons aligned to the peak of theta improves encoding, whereas stimulation aligned to the trough improves retrieval (Siegle and Wilson, 2014).

In rodents, theta-modulated gamma band activity has also distinguished encoding from retrieval (Bragin et al., 1995; Colgin, 2015). Distinct slow (~30 to 50 Hz) and mid (~50 Hz to 100 Hz) gamma oscillations are observed and generated by separate neural circuits, with maximal amplitudes at the peak and trough of the theta rhythm, respectively (Colgin, 2016). Recent work has also identified unique theta-nested gamma oscillations observed within individual theta cycles (Lopes-Dos-Santos et al., 2018), providing support for the notion that fast and slow gamma separately mediate encoding of novel information and memory retrieval. Intracranial recordings in humans have identified ripple oscillations (~80 to 100 Hz) that are involved in memory retrieval and consolidation (Axmacher et al., 2008; Staresina et al., 2015; Vaz et al., 2019) and exhibit phase amplitude coupling (PAC) with hippocampal theta phase (Staresina et al., 2015). Similar ripple oscillations are also prevalent in nonhuman primates during visual search (Leonard and Hoffman, 2017; Leonard et al., 2015), raising the possibility that they play an active role in exploration. Because evidence for memory-related hippocampal theta to gamma PAC in humans has primarily focused on verbal learning tasks (Lega et al., 2016; Mormann et al., 2005), it is not known how changes in PAC during exploration translate from animal models to similar behaviors in humans.

Here, we recorded eye movements and intracranial hippocampal recordings as neurosurgical patients performed an associative spatial memory task. We hypothesized that theta oscillations would modulate distinct gamma frequencies when eye movements were driven by either associative novelty or retrieval. Indeed, previous work in humans has examined the relation between theta oscillations and memory in verbal recall tasks (Kahana et al., 2001; Lega et al., 2012; Sederberg et al., 2003), including theta-gamma phase amplitude coupling (Lega et al., 2016; Mormann et al., 2005; Vaz et al., 2017). These studies examined encoding and retrieval in isolated task epochs, raising the question of how theta supports these memory processes when they rapidly co-occur. In our spatial memory task, subjects encountered previously studied objects in either their original or updated spatial locations. When an object appeared in an updated location, subjects directed viewing to both the updated and original locations iteratively over the course of the trial. Simultaneous acquisition of eye movements and intracranial EEG allowed us to relate theta to the timing of encoding and retrieval processes with exceptional temporal resolution. In doing so, we systematically tested the hypothesis that hippocampal theta influences when different memory processes occur during exploratory viewing. If the strength of encoding and retrieval are modulated by theta phase, fixations driven by each process should be phase-locked to the theta rhythm. In addition, eye-movements tied to distinct encoding and retrieval processes should be accompanied by distinct states of hippocampal PAC. As such, this experiment determines how the hippocampus contributes to learning and coordinates dynamic encoding and retrieval operations during visual exploration in humans.

## Results

### Direct brain recordings linked to memory-driven eye movements

Patients performed an associative spatial memory task (Fig. 1a), consisting of three phases, while we simultaneously acquired recordings of eye movements and local field potentials from the hippocampus (Fig. 1b). During the study phase, patients learned the spatial location of 16 objects presented sequentially on a background scene. Next, during a refresh phase, objects were represented in either their original (Match) or updated (Mismatch) spatial locations, with two visual cues (small red dots) indicating potential alternate locations. One visual cue always indicated the object’s original location during Mismatch trials. After viewing each stimulus, patients determined via button press whether each object was presented in its original or updated location. During a final recognition phase, patients viewed each object in three locations and attempted to identify the object’s original location.

**Figure 1.**
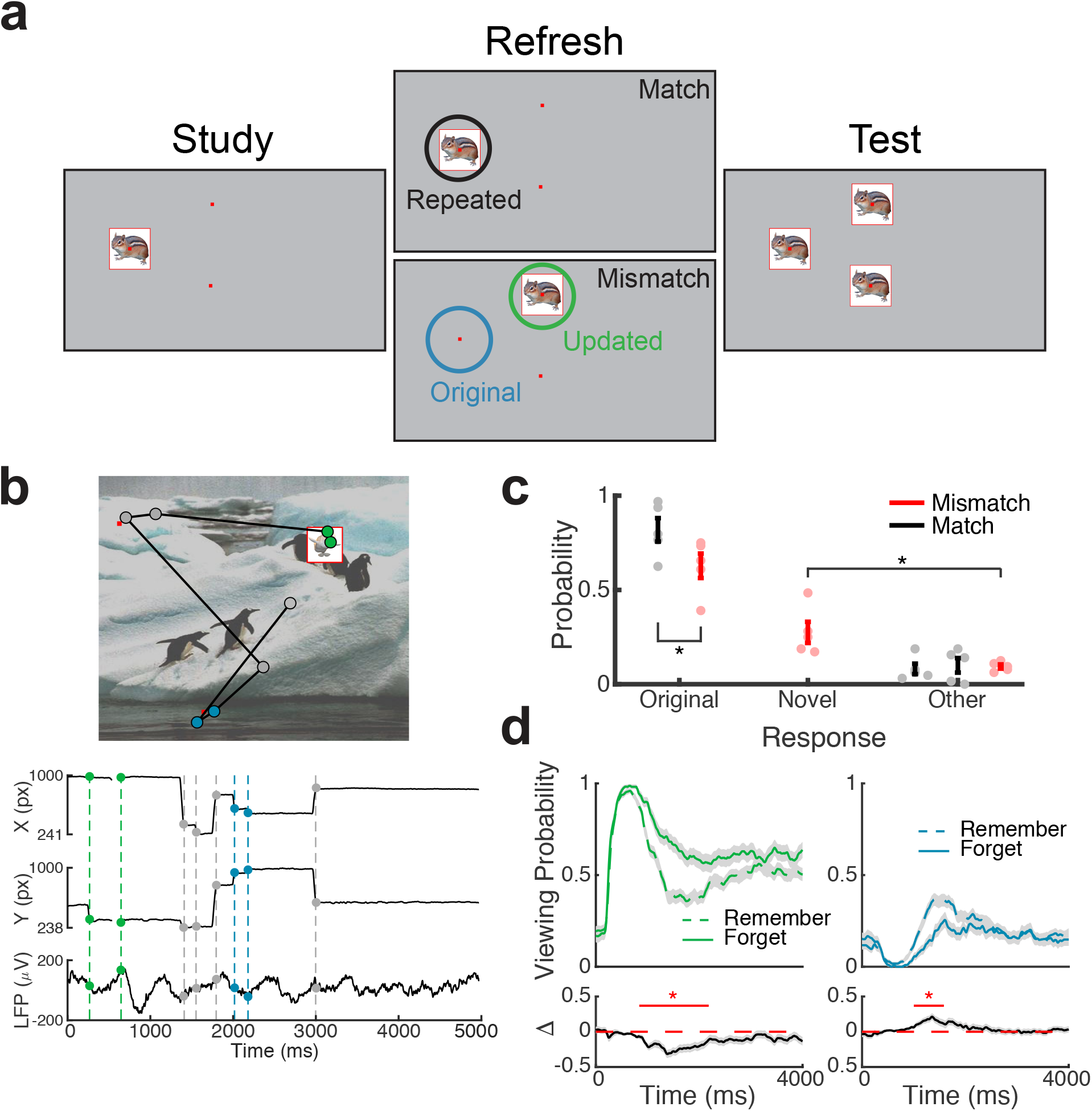
Direct brain recordings during memory-driven eye movements. (**a**) Spatial memory task. Example stimuli presented during each phase of the task. Viewing regions of interest (ROIs) for each trial type are indicated by circles on the Refresh phase. (**b**) Simultaneous recording of gaze position and hippocampal field potential during an example trial. Above, viewing scan path overlaid on the stimulus display for a Mismatch trial. Below, gaze position and concurrent signal for a bipolar pair in the hippocampus. The onset of fixations to viewing ROIs are denoted by colored circles. (**c**) Behavioral performance. Response proportions on the final recognition test for each viewing condition. Each point denotes a subject average; lines denote one SEM. (**d**) Viewing behavior on Mismatch trials predicts memory outcomes. The probability of viewing the novel (left) or original (right) object-location was compared on trials in which the original location was subsequently remembered or forgotten. Below, a subsequent memory effect was computed as the difference in viewing probability. Shaded areas depict ±SEM. Lines depict significant clusters (*P*_FWE_ < 0.05).

During the refresh phase, subjects were encouraged to visually explore the three cued locations to help inform their memory decision. Our primary analyses focused on the interplay of associative novelty and retrieval processes during these Mismatch trials, by linking hippocampal activity to eye movements directed to the original and updated locations. This approach enabled us to identify distinct hippocampal mechanisms linked to these cognitive processes. In addition, we evaluated the impact of viewing behaviors and electrophysiological states on final recognition performance.

We measured overall task performance by computing accuracy on the final recognition test. Patients performed the task well, with the original object location remembered on 82 (± 6 SEM) percent of recognition trials (Fig. 1c). Recognition accuracy was significantly impaired on Mismatch relative to Match trials (paired t-test, *t*_4_= 9.83, *P* = 0.0006). To confirm that eye movements during the refresh phase were tied to memory processes, we examined the changes in viewing behaviors and memory outcomes on the final recognition test (Table 1). On average, participants made more fixations to the presented object during Match (paired t-test, *t*_4_= 12.9, *P* = 0.0002) and Mismatch (paired t-test, *t*_4_ = 17.7, *P* = 0.0001) trials than to the other two cued locations. Notably, the number of fixations to the object was reduced on Mismatch relative to Match trials (paired t-test, *t*_4_ = −8.6, *P* = 0.0009), indicating that subjects explored additional visual features during Mismatch. In addition, fixation durations to objects were longer than matched spatial cues (paired t-test, all *t*_4_ > 3.39, *P*s < 0.023) and were comparable between Match and Mismatch trials (paired t-test, *t*_4_ = 1.49, *P* = 0.21).

**Table 1.** Task-related eye movement behavior.

|  | Mismatch |  |  | Match |  |  |
| --- | --- | --- | --- | --- | --- | --- |
|  | Original | Updated | Other | Repeated | Other | Other |
| Fixations per trial (N) | 2.4 (0.3) | 4.6 (0.4) | 1.4 (0.3) | 5.3 (0.4) | 1.7 (0.4) | 1.4 (0.3) |
| Count SME (*t*) | 1.4 | -4.6* | -0.3 | 3.7* | -0.4 | -1.7 |
| Fixation duration (ms) | 210 (11) | 388 (59) | 212 (14) | 342 (39) | 199 (11) | 206 (6) |
| Duration SME (*t*) | 0.5 | 1.1 | 1.4 | -1.6 | -2.4 | -.6 |

\* $p < 0.05$ , parenthesis denote SEM

**Table 2.** Subject demographics.

|  | **S1** | **S2** | **S3** | **S4** | **S5** |
| --- | --- | --- | --- | --- | --- |
| **Age (years)** | 20 | 34 | 53 | 25 | 44 |
| **Sex** | M | M | M | F | F |
| **Full-scale IQ** | 94 | 109 | 105 | 121 | 92 |
| **Implanted Hemisphere** | Left | Left | Right | Left | Left |
| **Epileptic Focus** | Basal temporal | Basal temporal | Middle hippocampus | Basal temporal | Amygdala |
| **Etiology** | Cortical dysplasia | Dysembryoplastic neuroepithelial tumor | Focal cortical dysplasia | Low grade glioma | Mesial temporal sclerosis |
| **Duration of epilepsy (years)** | 10 | 10 | 8 | 3 | 41 |
| **Hippocampal contacts (n)** | 8 | 4 | 6 | 7 | 6 |

Viewing behavior on Match and Mismatch trials predicted final recognition performance. On Match trials the number of fixations to the repeated object predicted better memory for the original location, whereas the number of fixations to the updated location on Mismatch trials predicted memory updating (see Table 1 for details). To break down the timing of these memory-guided eye movements, we examined the proportion of time spent viewing each ROI across trials (Fig. 1d). We found that viewing preferences on Mismatch trials, but not Match trials, predicted later memory for the original object-locations. Following initial visual orienting to the novel stimulus, prolonged viewing of the novel object-location (738 to 2188 ms after object presentation) was associated with memory updating (nonparametric cluster test, *P*_FWE_ < 0.05). Viewing the original object-location during this time period (from 998 to 1584 ms) led to better memory for the original location (nonparametric cluster test, *P*_FWE_ < 0.05). Proportion of viewing over time during Match trials was not a significant predictor of final memory performance (*P*_FWE_ > 0.05). These findings suggest that interplay between memory processes and visual sampling during Mismatch trials determined whether memory updating would occur.

### Theta dependence of memory-guided eye movements

We analyzed direct recordings from 32 electrodes from 5 patients (Fig. 2a), focusing on depth electrodes in the hippocampus. We measured spectral power and phase from 1 Hz to 10 Hz, which includes both theta and low-theta (Jacobs, 2014; Watrous et al., 2013a) frequency bands previously associated with visual exploration and memory encoding (Jutras et al., 2013; Lega et al., 2012). We linked measures of spectral power and phase to individual fixation events to identify hippocampal states reflecting retrieval and novelty detection. We reasoned that hippocampal signaling prior to fixations would reflect a memory-guided initiation of the upcoming eye movement, whereas signaling following fixations would reflect a memory-based reaction to the visual input.

**Figure 2.**
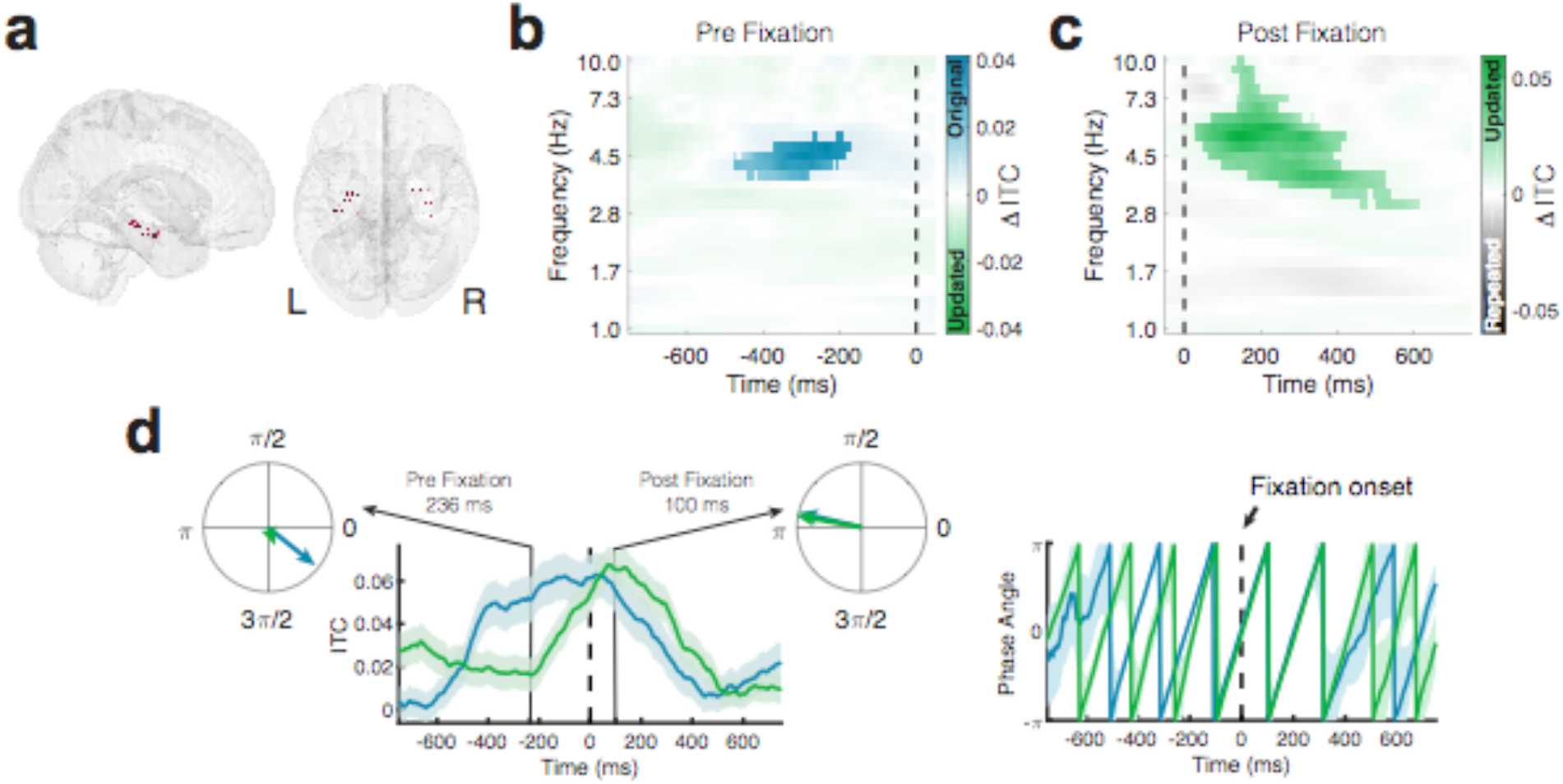
Phase-locking of memory-dependent eye movements to hippocampal theta. **(a)** Location of hippocampal electrodes in MNI space. **(b)** Increased phase locking precedes retrieval-dependent fixations. Significant differences (cluster *P*_FWE_ <0.05) in inter-trial phase clustering (ITC) between fixations (indicated by the dashed line) to original vs. updated object-locations on Mismatch trials are highlighted. See Figure S1 for related changes in theta power. **(c)** Novelty related modulations in hippocampal phase. Significant differences in ITC following fixations to the updated object-location on Mismatch trials and the repeated object-location on Match trials. **(d)** Left, timecourse of ITC for phase at 5 Hz. Polar plots depict the magnitude of ITC and preferred phase for fixations to either the original or updated object-locations on Mismatch trials. Right, alignment of phase angles prior to fixations on Mismatch trials. Shaded regions depict ±SEM.

To assess whether retrieval-guided eye movements during Mismatch trials occurred at specific phases of hippocampal oscillations, we compared the consistency of phase angles in the moments leading up to fixations to the original and updated locations using inter-trial phase coherence (ITC; Fig. 2b). We observed significantly greater phase-locking around 5 Hz prior to fixations to the original compared to the updated object-location on Mismatch trials (*P*_FWE_ < 0.05, nonparametric cluster correction). Importantly, the lack of theta phase-locking preceding fixations to the updated location could not be accounted for by decreased power of hippocampal theta oscillations (Fig. S1).

As fixations to the original object-location were frequently preceded by novelty-driven fixations (Fig. 1d), it is possible that the observed phase-locking effect resulted from novelty detection rather than retrieval. Two control analyses suggested this was not the case. First, ITC measured during fixations to the updated location did not differ depending on the target of the next saccade (i.e., either to the original or updated location; all *P*_FWE_ > 0.05). Second, we observed significantly increased theta phase-locking across subjects (one sample t-test, *t*_4_ = 2.95, *P* = 0.04) during the same time interval when we excluded retrieval-guided fixations that were preceded by fixations to the updated object-location (and may be confounded by novelty detection). Taken together, these results implicate hippocampal theta oscillations in guiding eye movements via memory retrieval.

We next asked whether theta phase was modulated following fixation onset. If theta phase is generally modulated by fixations during memory updating (i.e., during Mismatch trials), consistent phase-locking could occur irrespective of the viewing location. To test this possibility, we contrasted ITC between each type of fixation (to the original or updated location) on Mismatch trials with fixations to the repeated location on Match trials. We observed significantly (*P*_FWE_ < 0.05, nonparametric cluster correction) greater phase clustering following fixations to updated compared to repeated object-locations (Fig. 2c). We only observed this post-saccade phase-locking effect following fixations to updated locations, as ITC did not significantly differ between fixations to the Mismatch-original and Match-repeated locations. In addition, we did not find any significant differences between phase-locking during fixations to original and updated locations during Mismatch trials. The time course of theta clustering (Fig. 2d) reveals the timing of hippocampal retrieval and novelty processes; a pre-fixation retrieval effect and a reset of theta phase due to fixations to updated object-locations.

We next determined whether theta phase dynamics predicted memory performance on the final recognition test. We found that phase-locking following fixations to updated locations during Mismatch trials predicted subsequent memory performance (Fig. 3). Greater ITC at frequencies ranging from 3 to 10 Hz following fixations to the updated location was associated with better memory stability on the final recognition test. The ability of hippocampal phase-locking to predict memory outcomes was specific to viewing updated object-locations; we observed no significant subsequent memory effects following fixations to either original or repeated object-locations. Furthermore, these differences could not be accounted for by differences in power (all *P*_FWE_ > 0.05, nonparametric cluster correction; data not shown). Taken together, these results demonstrate that hippocampal theta oscillations are tightly coupled to distinct viewing behaviors driven by associative novelty and memory retrieval.

**Figure 3.**
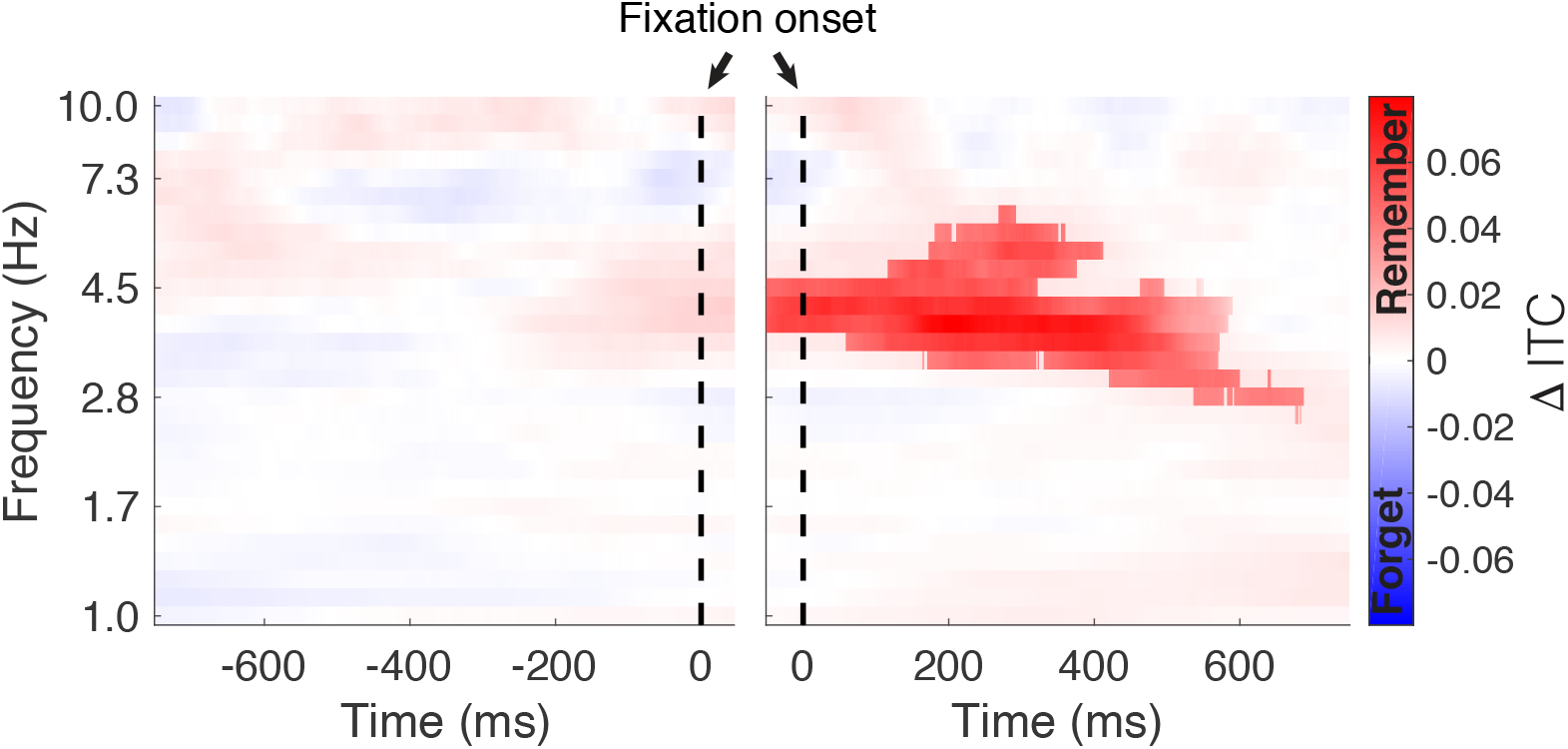
Phase locking of hippocampal theta predicts subsequent memory. Phase-locking of hippocampal theta predicts subsequent memory. Time-frequency plots depict differences is inter-trial phase coherence (ITC) between fixations to subsequently remembered and forgotten Mismatch trials. Significant (*P*_FWE_ < 0.05, nonparametric cluster corrected) increases in phase-locking were observed following the fixation onset.

### Theta to gamma phase amplitude coupling predicts memory updating

Having identified a consistent relationship between theta phase and specific viewing behaviors, we examined the relationship between theta phase and the amplitude of high frequency (80-200 Hz) gamma band activity. Phase-amplitude coupling (PAC) between theta and gamma has been proposed as a mechanism for separating memory representations (Hasselmo and Eichenbaum, 2005), with supporting evidence in both animal models (Tort et al., 2009) and humans (Axmacher et al., 2010; Heusser et al., 2016; Lega et al., 2016; Lisman and Jensen, 2013; Vaz et al., 2017). We used the modulation index (MI) to quantify PAC between the phase of theta (ranging from 1 to 10 Hz) and gamma amplitude (Tort et al., 2010).

We focused our PAC analyses on two comparisons of interest: associative novelty and memory retrieval. Results from an example electrode are depicted in Figure 4, showing increases in PAC related to associative novelty. For a given bipolar recording pair (Fig. 4a), we computed gamma amplitude as a function of theta phase during each trial type of interest. These measures were used to compute the MI, which measures PAC as the difference between the observed amplitude distribution and a uniform distribution (Fig. 4b, dashed line). To make sure observed differences in PAC did not result from non-stationarities in the data or common task-evoked changes in amplitude and phase (Aru et al., 2015), we permuted phase information across trials for each condition and computed a normalized score (MI_Z_) based on this null distribution. We assessed changes in MI_Z_ across conditions (Fig. 4c) to identify memory-related changes in PAC. Across subjects, we found 25% of hippocampal electrodes exhibited significant differences in PAC driven by associative novelty (i.e., differences between fixations to updated or repeated locations), significantly more than expected by chance (*P* < 0.001, binomial test). Only 9% of electrodes exhibited significant differences in PAC preceding fixations to the original versus updated locations on Mismatch trials (*P* = 0.14, binomial test).

**Figure 4.**
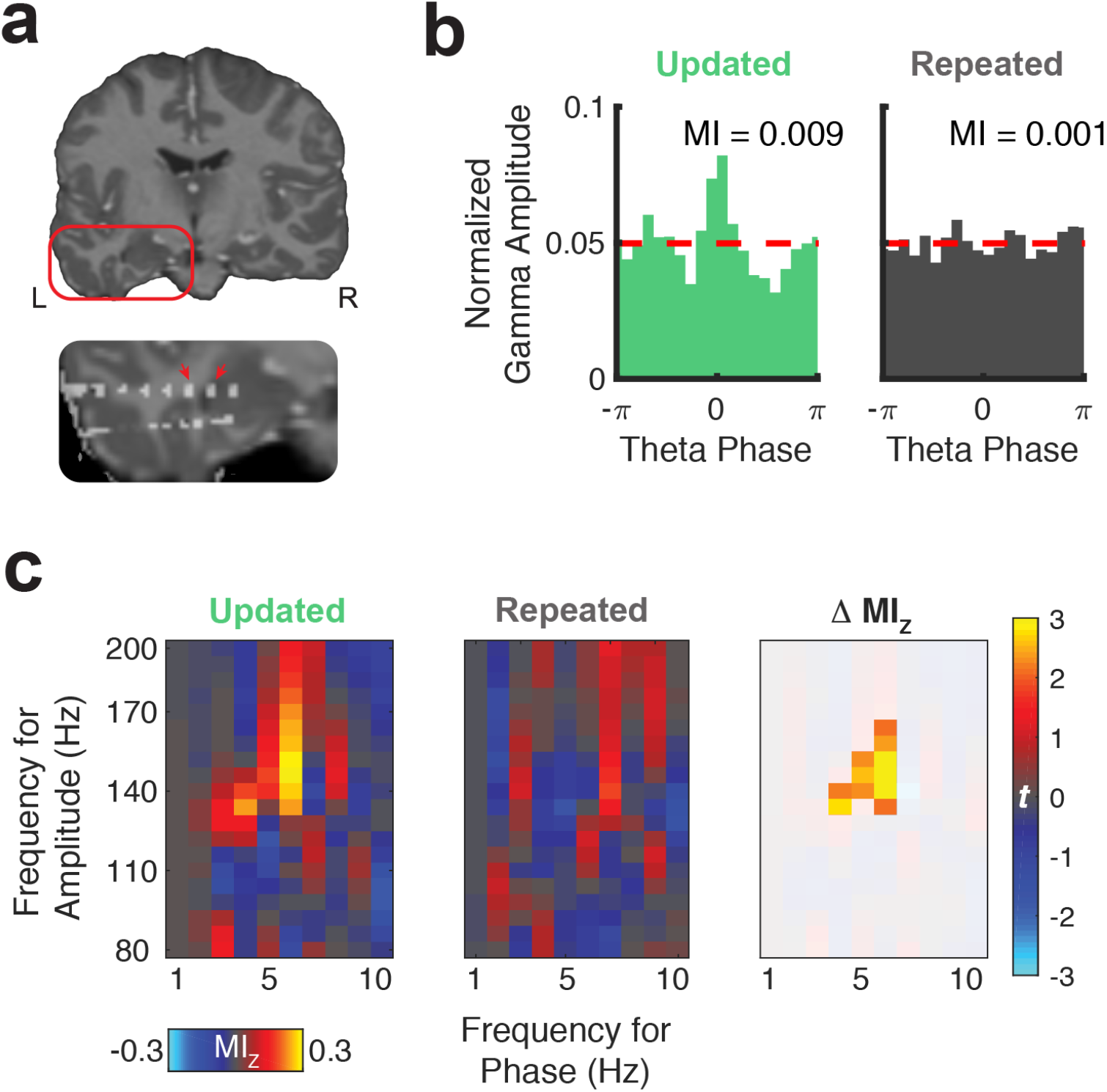
Representative theta to gamma phase amplitude coupling at an individual electrode. (**a**) Re-referenced bipolar recording from contacts in the left hippocampus and adjacent white matter. (**b**) Normalized amplitude distributions reveal novelty-related modulation of gamma (140 Hz) amplitude by theta (6 Hz) phase at this recording site. MI, modulation index. Dashed line denotes normalized gamma amplitude under a uniform distribution. (**c**) Left, comodulograms depict increased PAC (z-scored MI, constructed from trial-shuffled surrogate data) during fixations to updated compared to repeated object-locations. Right, the statistical map depicts a cluster of significant (*P*_FWE_ < 0.05, nonparametric cluster corrected) cross-frequency interactions.

Next, we tested for group-level differences in PAC during specific viewing behaviors. To account for variability in theta frequency across subjects and electrodes, we selected the theta frequency that exhibited the greatest magnitude MI_Z_ from 4 to 6 Hz, irrespective of condition. We first examined if theta to gamma PAC was sensitive to associative novelty by contrasting PAC following fixations to updated versus repeated objects. We found significantly increased theta to gamma (80 to 100 Hz) PAC during fixations to updated objects, indicating that gamma amplitude was more dependent on theta phase when visual stimuli conflicted with memory (Fig. 5a).

**Figure 5.**
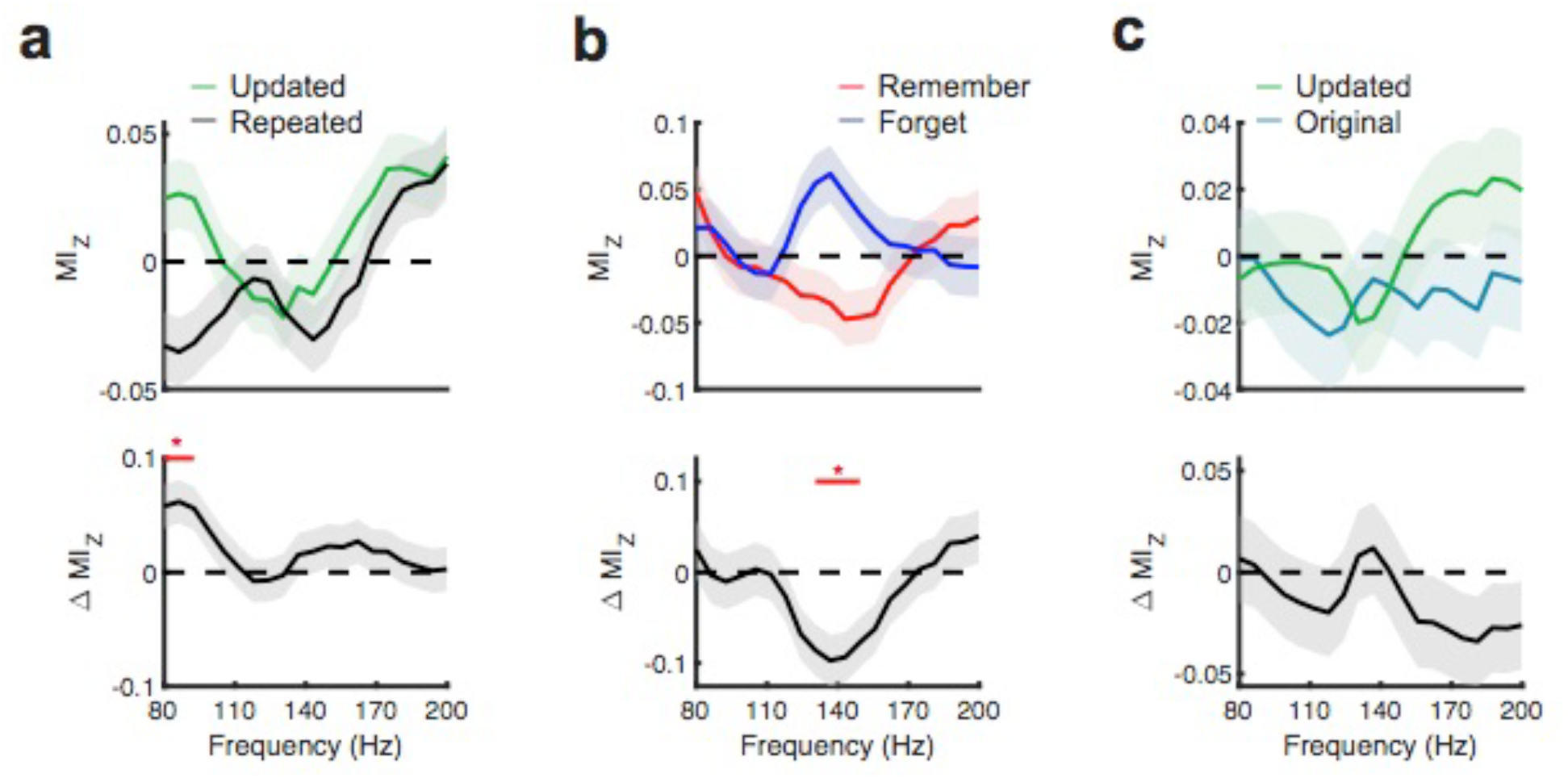
Hippocampal phase amplitude coupling predicts novelty detection and memory updating. (**a**) Top, post-saccade changes in PAC during fixations to updated and repeated object-locations are displayed for a range of gamma amplitudes. Below, significant (*P*_FWE_< 0.05, nonparametric cluster corrected) increases in PAC during related to novelty are indicated. (**b**) Theta to gamma PAC during object fixations on Mismatch trials varies with memory outcome. A significant (*P*_FWE_ < 0.05, nonparametric cluster corrected), negative subsequent memory effects is depicted in the bottom panel. (**c**) PAC did not differ in the moments leading up to fixations to updated and original object-locations during Mismatch trials.

Given significant PAC effects during updated fixations, we evaluated whether PAC during these fixations predicted subsequent memory performance. We found that increases in theta to high-gamma (130 to 150 Hz) PAC were significantly greater on trials where the original object location was subsequently forgotten (Fig. 5b), suggesting that increased theta-high gamma PAC corresponded to enhanced encoding of the updated location. Consistent with electrode-level analyses, we did not observe a consistent relationship between PAC and retrieval-related eye movements (Fig. 5c). We examined the specificity of these effects to different frequency ranges by repeating this analysis for low-theta (1-3 Hz) and faster theta/alpha frequencies (7-10 Hz). We did not observe significant differences in PAC in these ranges, suggesting specificity of memory-related PAC in the 4-6 Hz theta range. The observed differences in theta to gamma PAC reflect different processing states in hippocampal networks, wherein asynchronous local activity is necessary to segregate novel perceptual information from previously encoded memories.

## Discussion

We set out to examine the relationship between hippocampal theta oscillations and memory-driven viewing behaviors during visual exploration. To achieve this goal, we examined simultaneously recorded hippocampal potentials and eye movements while neurosurgical patients performed a spatial memory task. This paradigm allowed the disambiguation of eye movements driven by associative novelty (i.e., objects presented in updated spatial locations) and memory retrieval. We discovered that these two distinct viewing behaviors were uniquely tied to theta: phase locking preceded retrieval-guided eye movements and followed novelty-driven fixations, predicting memory performance. Analysis of theta to gamma PAC confirmed a potential mechanism by which the theta rhythm supports both encoding and retrieval. Modulation of gamma (80-100 Hz) amplitude by the theta rhythm increased during detection of updated object locations. Further, increased theta to high-gamma PAC predicted memory updating, as determined by a subsequent memory test. These data support theories of active hippocampal involvement in visual exploration (Voss et al., 2017) and provide novel evidence that theta oscillations support both retrieval-and novelty-dependent viewing behaviors.

Based upon electrophysiological studies in humans and primates (Hoffman et al., 2013; Jutras et al., 2013) it has become apparent that hippocampal memory representations are used to guide saccades to behaviorally relevant locations (Meister and Buffalo, 2016). Of particular relevance to the present work, visual exploration of novel, but not repeated, scenes leads to a reset of hippocampal theta oscillations (Jutras et al., 2013). The consistency of this hippocampal phase reset predicts the success of novel encoding. In the present work, we provide evidence for conserved hippocampal processing in humans, by demonstrating increased phase consistency of theta oscillations following fixations to novel locations, which were further associated with improved memory performance. Recent work using fMRI (Liu et al., 2017) also suggests that visual exploration of scenes is associated with hippocampal function, as the number of fixations during encoding predicted both hippocampal activity and subsequent memory. This relationship between hippocampal activity and viewing behavior was limited to novel stimuli (but not repeated stimuli), suggesting hippocampal activity and exploratory eye movements are linked specifically when encoding novel content. Using our spatial memory paradigm to identify retrieval-dependent eye movements, we found theta phase-locking occurred prior to saccade onset. Taken together with previous studies that have tied hippocampal activity to retrieval-guided eye movements (Bridge et al., 2017; Hannula and Ranganath, 2009) these results provide evidence that visual exploration is dependent upon the interplay of separate retrieval and novelty-detection mechanisms that underpin learning.

With the high temporal resolution eye movements and hippocampal potentials, our data are uniquely suited to clarify the hippocampal mechanisms that drive learning under associative novelty. Notably, we found increased theta phase-locking and power following fixations to mismatched spatial locations, consistent with previous hippocampal recordings in humans (Chen et al., 2013). The consistent timing of these effects following fixation onset provide further evidence that hippocampal theta is involved in the computation of mismatch signals, as opposed to providing signals of perceptual familiarity that could guide viewing to novel locations. This extends previous fMRI work which has established the involvement of the hippocampus in associative novelty detection (Kumaran and Maguire, 2006, 2007a) and binding (Bridge and Voss, 2014b), specifically supporting the idea that the hippocampus acts as a comparator between new sensory inputs and prior memory (Kumaran and Maguire, 2007b; Lisman, 1999; Lisman and Grace, 2005; Vinogradova, 2001).

The observed relationship between the phase of theta oscillations and gamma amplitude provides insight into the mechanism by which hippocampal theta can promote learning under associative novelty. Individual theta cycles have been proposed to segregate neuronal activity into functional packets based on coactivation of neuronal populations during distinct time windows (Buzsaki and Moser, 2013; Hasselmo et al., 2002; Renno-Costa and Tort, 2017). Studies of rodent electrophysiology within the hippocampus have revealed slow (~30 to 50 Hz) and fast (~50 to 100 Hz) gamma oscillations within area CA1 that exhibit PAC to the theta rhythm, with distinct preferred phases (Colgin et al., 2009; Yamamoto et al., 2014). Based on this evidence, it has been proposed that distinct phases of the theta rhythm in CA1 reflect different network states that differentially support memory encoding and retrieval (Colgin, 2015).

We found a relationship between theta phase and gamma amplitude during fixations to updated stimulus locations. Consistent with previous investigations of PAC and memory processing (Lega et al., 2016; Vaz et al., 2017), increased PAC was associated with impaired memory function. Inspection of raw traces (e.g., Fig. 1b) revealed that the observed differences in PAC resulted from nested oscillations, as opposed to the modulation in the amplitude of sharp waveforms. Distinct neurophysiological states, defined by theta phase and modulation of separate gamma bands, therefore direct viewing from memory as opposed to novel information in the environment. While we did not observe theta-dependent modulation of gamma amplitude during retrieval, this could result from a number of methodological constrains, such as biased sampling to the anterior hippocampus and potential obscuring of narrowband gamma signals commonly observed in microelectrodes due to summation across larger diameter electrodes.

Despite the emerging consensus in the rodent that theta oscillations in the hippocampus are responsible for segregating neuronal computations involved in encoding and retrieval operations (Colgin, 2015, 2016; Hasselmo et al., 2002), translating these findings to observations in humans has proven challenging. Comparative studies between rodents and humans suggest that theta rhythms in the human are commonly observed at lower frequencies (3 vs. 8 Hz) during exploratory behaviors (Watrous et al., 2013a). Additionally, studies have demonstrated multiple sources of theta rhythms in the medial temporal lobe (Lega et al., 2012; Mormann et al., 2008), including a “low-theta” or delta band (1-4 Hz) in addition to the typical theta band (4-8 Hz). Based on observations of encoding-related increases in power (Burke et al., 2013; Lega et al., 2012; Miller et al., 2018) and PAC during verbal encoding are specific to the "low-theta" band (Lega et al., 2016), it has been proposed that lower frequency theta in humans reflects a homologue of rodent theta (Jacobs, 2014). In contrast to this body of work, we observed faster (i.e., predominantly greater than 4 Hz) memory-related theta dynamics. We believe this difference stems from the emphasis on visual information in our task, resulting in a task-dependence in the frequency of theta oscillations. Higher frequency theta effects have been observed during visual search (Hoffman et al., 2013), in which alignment between saccades and theta oscillations was focused between 6-8 Hz. Our findings of novelty effects in the 4-10 Hz range are consistent with work in nonhuman primates (Jutras et al., 2013), which demonstrated that saccades during visual exploration caused resets in hippocampal oscillations predominantly in the 8-11 Hz range. As in the present study, the magnitude of these resets predicted the success of encoding during memory formation. Multiple factors could account for changes in the speed of hippocampal theta, including the type of representation being processed in the medial temporal lobe (Watrous et al., 2013b), spatial attention required by a given task, or the rate of fixations. Future studies are necessary to determine what factors determine the speed of hippocampal theta oscillations and their relevance to different forms of memory.

While our analysis of theta oscillations was restricted to electrodes in the hippocampus, memory-guided exploratory behaviors depend upon interactions between distributed cortical systems (Voss et al., 2017), particularly those involved in the representing features within a scene, the spatial relationships between these features, and transforming these memory representations into an oculomotor plan based on current visual input. Kilian et al. (2012) discovered cells within the entorhinal cortex of macaques that coded the location of fixations in a grid-like fashion during a free-viewing task, serving as a potential mechanism to provide a scale-invariant representation of fixation locations within the scene (Bicanski and Burgess, 2019). Similar grid-like modulation of entorhinal activity has been observed in humans using fMRI (Julian et al., 2018; Nau et al., 2018), providing converging evidence across species that the entorhinal system may provide a spatial framework for memory-guided viewing. As such, synchronous theta oscillations between the hippocampus and entorhinal systems would provide the spatial coding necessary to inform the oculomotor system of memory-relevant information. Though we were limited by electrode coverage, examining entorhinal-hippocampal synchrony and interactions between the hippocampus and other cortical systems should be a key aim for future studies.

One potential caveat is that the observed eye movements in this study do no reflect natural exploratory behaviors per-se, but are rather driven by demands to learn and maintain the original object location throughout the task. If so, it is unclear how stereotyped these retrieval-dependent eye movements would be during unconstrained visual exploration. Free-viewing paradigms could build on this theoretical framework, and determine the extent to which hippocampal-dependent viewing behaviors occur without task constraints. Given the observed retrieval phase-locking effects occurred well before saccade initiation, it is likely that the hippocampus plays a causal role in generating these eye movements. Causal manipulation of hippocampal theta, including stimulation-based approaches could be used to test this hypothesis, coupled with growing evidence for disruptions in viewing behaviors from amnesic patients with hippocampal damage (Hannula et al., 2007; Lucas et al., 2019; Olsen et al., 2016; Ryan et al., 2000; Smith et al., 2006).

In conclusion, encoding and retrieval dependent eye movements are time locked to the phase of the hippocampal theta rhythm which supports the model that distinct phases of the theta cycle segregate neural processing of information related to the present and the past (Colgin, 2016; Hasselmo et al., 2002; Hasselmo and Eichenbaum, 2005). Distinct encoding and retrieval states at different phases of the theta cycle would allow for structured comparison of the present and the past. Akin to spatial attention shifting between multiple locations relevant to a task at hand (Landau et al., 2015; Re et al., 2019), hippocampal theta could coordinate visual sampling between novel content in the environment and memory-rich spatial locations. The hippocampus thereby contributes to memory-guided behaviors through coordinated sampling of current and past perceptual states.

## Acknowledgments

We are grateful to Drs. Christina Zelano and Joel Voss for helpful discussions and the Laboratory of Human Neuroscience at Northwestern University for sharing resources; D.J.B acknowledges the support of NIH/National Institute of Mental Health (NIMH; grant R21MH115366) and National Center for Advancing Translational Sciences, Grant Number UL1TR001422. This work was supported in part by a National Institute of Neurological Disorders and Stroke grant T32NS047987 to J.E.K.

## Author Contributions

Conceptualization, J.E.K. and D.J.B.; Formal Analysis, J.E.K. and D.J.B.; Investigation D.J.B., J.E.K., and A.S.N.; Resources, S.V.H., S.S., J.M.R. and J.W.T; Writing – Original Draft, J.E.K.; Writing – Review and Editing; D.J.B, J.E.K., and A.S.N.; Funding Acquisition, D.J.B.; Supervision, D.J.B.

## Methods

### Participants

5 patients (3 male; see Table 1 for demographic information) with refractory epilepsy performed our associative memory task during their stay at Northwestern Memorial Hospital (Chicago, IL). All patients had depth electrodes implanted in the hippocampus as part of neurosurgical monitoring prior to elective surgery. Written informed consent was acquired from all patients prior to participation in the research protocol in accordance with the Northwestern University Institutional Review Board.

### Experimental paradigm

We tested memory for associations between objects and their spatial locations using a novel spatial memory task. This task consisted of 3 distinct phases (Study, Refresh, Recognition), with each phase separated by a 60s distractor (presentation of unrelated visual stimuli). Subjects performed 8 blocks in which they learned spatial locations for a sequence of 16 objects. Eye movements were recorded during each phase of the task, with 5-point gaze calibration performed before each phase. Objects were 128 images of real-life objects from the Bank of Standardized Stimuli (Brodeur et al., 2010). During each phase of the task, objects were presented at 3º of visual angle, with a red square of 0.2º of visual angle centered on each object. Stimuli were presented on a 23.6” monitor with a 120 Hz refresh rate from a stimulus control laptop. Synchronization pulses were sent from the stimulus control laptop to the clinical recording system using a DAQ control board, allowing alignment of electrophysiological and behavioral data.

At the beginning of the Study phase, a unique background image appeared for 5s to allow familiarization with the scene that could serve as a reference for the spatial location of each object. Throughout the remainder of the Study phase, a sequence of 16 objects were presented at distinct locations superimposed on the background scene. At the start of each study trial, a fixation cross flashed twice on the screen (250 ms per flash, separated by 250 ms of the background scene) to cue the location of the next object. The fixation cross remained on the screen for a duration of 2s, followed by presentation of the object for 3s.

Next, subjects were tested on their spatial memory for each of the objects during the Refresh phase. During this phase of the task, three location cues indicated by small red squares (0.2º of visual angle) were presented in an equilateral triangle (randomly selected distance for each stimulus, mean distance of 12º and a range of 5.9-21.1º of visual angle across presented arrays). The object was presented at one of these three locations. Importantly, one of these locations was the object’s original location. On each block, half of the trials were randomly assigned to the Mismatch condition, in which the object was presented at one of the two novel locations. On the Match trials, the object was presented in its original location. Each trial began with the presentation of the background scene for 1s followed by a fixation cross at the center of the screen for 1s, at which point the object and location cues appeared for 5s. Following stimulus presentation, memory for original location of each item was tested. Subjects identified whether the object was in its original location, a new location, or if they were unsure by clicking a box that said: Same, Different, or Unsure.

Each block concluded with the Recognition phase which served as a final memory test for the original object locations. The background scene appeared for 1s, followed by the presentation of a fixation cross in the center of the screen for 1s. Then, each object was presented at all three locations for a duration of 5s. Following stimulus presentation, subjects selected the original object location using a three-button response pad.

### Eye tracking

Eye movements were recorded at 500 Hz using an Eyelink 1000 remote tracking system (SR Research, Ontario, Canada). Continuous eye-movement records were parsed into fixation, saccade, and blink events. Motion (0.15º), velocity (30º/s) and acceleration (8000º/s^2^) thresholds were used to identify saccade events. Blinks were identified based on pupil size, and remaining epochs below detection thresholds were classified as fixations. The location of each fixation event was computed as the average gaze position throughout the duration of the fixation. Circular viewing regions of interest (ROIs) were constructed based on a distance of 6º from one of the three potential object locations. We focused our analysis of hippocampal activity to the subset of fixation events greater than 80 ms in duration that occurred 500 ms after object presentation and 500 ms prior to the end of each trial to avoid stimulus onset and offset effects.

### Intracranial recordings

A combination of depth electrodes (Integra Life Sciences, Plainsboro NJ; AD-TECH Medical Instrument Co., Racine, WI; DIXI Medical, Besançon, France) as well as subdural grids and strips. Electrode spacing on hippocampal depths was 5mm. Electrophysiological data were recorded to a clinical reference using a Nihon Kohden amplifier with a sample rate of 1-2 kHz with a bandpass filter from 0.6 to 600 Hz. Data were re-referenced to a bipolar montage and downsampled to 500 Hz as part of preprocessing. Line noise was reduced by application of a band-stop 4^th^ order Butterworth filter. To rule out the possibility that epileptiform activity influenced our analyses, electrodes that exhibited inter-ictal spiking were excluded from analysis. In addition, all analyses were repeated after excluding contacts within the seizure onset zone (2 electrodes in S3). There observed results were qualitatively identical, with no statistical differences when including all electrodes (all *P >* 0.05).

### Anatomical localization

Post-implant CT (n=4) or T1 weighted structural images (n=1) were coregistered with presurgical T1 weighted structural MRIs using SPM12. Subdural electrodes were localized by reconstructing whole-brain cortical surfaces from pre-implant T1-weighted MRIs using the computational anatomy toolbox (Dahnke et al., 2013) and snapping electrode centroids to the cortical surface based on energy minimization (Dykstra et al., 2012). All T1-weighted MRI scans were normalized to MNI space by using a combination of affine and nonlinear registration steps, bias correction, and segmentation into grey matter, white matter, and cerebrospinal fluid components. Deformations from the normalization procedure were applied to individual electrode locations identified on post-implant CT images or structural images using Bioimage Suite (https://medicine.yale.edu/bioimaging/suite/).

### Spectral decomposition

To examine oscillatory processes in the hippocampus, we decomposed bipolar recordings into measures of spectral phase and power using the continuous Morlet wavelet transform (wave number 5) across 30 logarithmically spaced frequencies from 1 to 10 Hz. We examined 1500 ms windows surrounding each fixation event of interest, with a 1250 ms buffer to prevent edge artifacts.

### Phase-locking analyses

We examined the relationship between the hippocampal theta rhythm and individual eye movements by computing the inter-trial phase coherence (a measure of phase-locking):

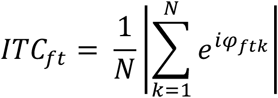

for a given time (*t*) and frequency (*f*), where *N* is the total number of individual trials, *k*, and *e*^*iφ*^ is the polar representation of the phase angle, *φ*. This measure was computed separately for individual conditions of interest (e.g., fixations to a specific region of interest on the display). As this measure is biased by the number of observations, with fewer observations leading to inflated ITC measures, we used a random subsampling approach to ensure that the number of observations were matched prior to statistical testing.

### Phase amplitude coupling analyses

Cross-frequency coupling between the phase of theta and gamma amplitude was computed using MI (Tort et al., 2010). MI is defined as the deviation in an amplitude distribution (across phases) from a uniform distribution, an adaptation from Kullback-Leibler distance (Kullback and Leibler, 1951), *D*_*KL*_, that normalizes the range of the distance between zero and one:

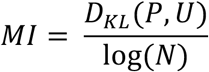

where *P* is the normalized amplitude distribution as a function of phase, *U* is a uniform distribution, and *N* is the number of phase bins. For all presented analyses, we used 20 phase bins of 18º. MI takes values greater than zero when the observed amplitude varies with phase and is equal to zero when the distribution is uniform.

To circumvent the relatively short epochs in which we analyzed cross-frequency coupling (constrained by the frequency of eye movements during our task), we computed a standardized measure of the modulation index, MI_Z_, via a surrogate control analysis. Specifically, for each trial and frequency combination, we permuted the observed phase timeseries across trials (separately for each condition of interest). This procedure was repeated 1000 times, resulting in a null distribution of MI values that could be explained by random (or condition-evoked) variations in the observed signal rather than true coupling between theta phase and gamma amplitude. MI_Z_ was measured as the difference between the observed MI and mean of the surrogate distribution, in units of standard deviations. These measures were used for all subsequent analysis of cross-frequency coupling.

### Statistical analyses

We adopted a nonparametric permutation-based approach (Maris and Oostenveld, 2007) to correct for multiple comparisons across time and frequencies. When comparing differences in ITC or power between different types of fixations, we constructed a null distribution of differences by permuting the assignment of condition labels, blocked at the subject level. This null distribution was used to define an independent cluster-forming threshold for each observed measure (e.g., ITC at a specific time-frequency pair). When measures would be biased by the number of observations per condition (e.g., differences in ITC), random subsampling was used to equate the number of observations per condition. Individual clusters were considered significant (*P*_FWE_ < 0.05) if the summed statistic within each observed cluster exceeded 95% or 97.5% of clusters in the null distribution for one-and two-tailed tests, respectively. For tests comparing the relationship between the phase of an oscillation and spectral power, null distributions were constructed by permuting the phase timeseries across trials within each condition, per electrode and subject. These null distributions were used to standardize measures of phase amplitude coupling prior to statistical testing, as described above.

## Supplemental Information

**Figure S1.**
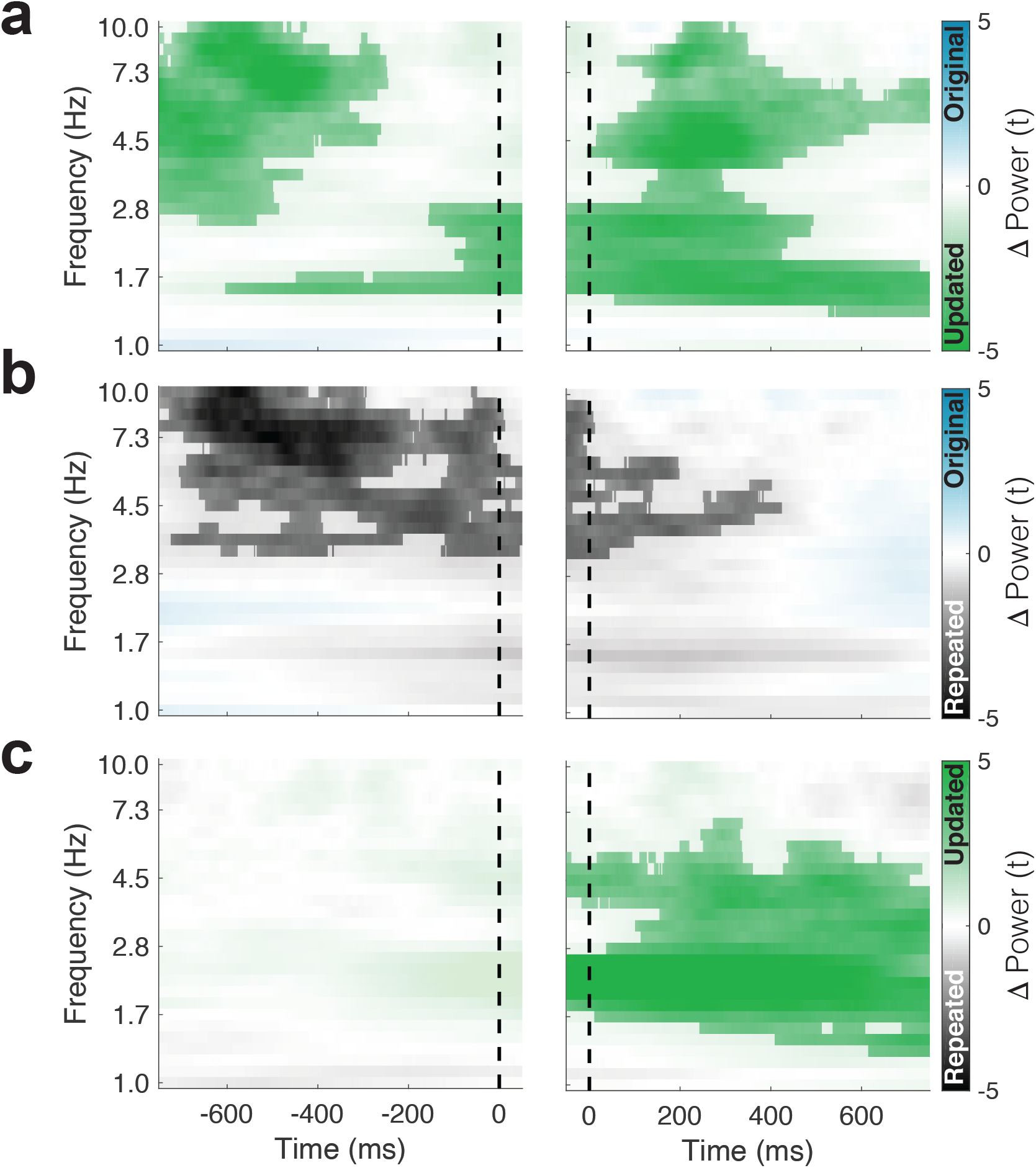
Eye-movement related changes in theta power (related to Figure 2). (**a**) Increased theta power precedes (left) and follow (right) fixations to the updated compared to the original object-location. (**b**) Same as **a**, but a contrast of theta power during fixations to the original object-location on Mismatch trials to the repeated object-location on Match trials. (**c**) Same as in **a**, but focusing on novelty-related changes in power. Increased low frequency power is identified following fixations to updated versus repeated object-locations. The vertical dashed line indicates the time of fixation onset. Significant clusters (*P*_FWE_ < 0.05, nonparametric cluster correction) are highlighted.

